# InterMap: Accelerated Detection of Interaction Fingerprints on Large-Scale Molecular Ensembles

**DOI:** 10.64898/2025.12.15.694195

**Authors:** Ernesto Fajardo-Díaz, Emmanuelle Bignon, François Dehez, Yasaman Karami, Roy González-Alemán

## Abstract

**Motivation:** Molecular dynamics is a key technique for exploring biomolecular systems at the atomic level. The rapid growth in accessible system sizes and timescales has intensified the need for efficient post-processing methods that extract meaningful insights from the resulting data. Interaction fingerprint (IFP) analyses are a valuable tool for elucidating key atomic interactions within molecular ensembles, yet current specialized software often struggle with extensive trajectories or complex systems. Here, we introduce InterMap, a Python package designed to accelerate IFP detection on large-scale molecular ensembles.

**Results:** By actively exploiting k-d trees, InterMap efficiently handles the massive amount of distance calculations necessary to detect IFPs, particularly when dealing with intra-molecular interactions. The seamless integration with MDAnalysis ensures broad format compatibility and allows using SMARTS patterns for flexible interaction definitions. InterMap adopts a deeply compressed binary encoding to manage IFPs, which makes it very memory-friendly. Furthermore, convenient interactive visualizations are provided to enhance data interpretation through a locally hosted web-browser application. Benchmark results indicate that InterMap significantly outperforms existing tools for processing complex biomolecular systems, achieving up to a 99% reduction in both runtime and peak memory usage.

**Availability:** InterMap’s code and issue tracker are available at https://github.com/Delta-Research-Team/intermap.git, while documentation and tutorials can be found at https://delta-research-team.github.io/intermap/.

**Supplementary information:** Supplementary data are available at *Nucleic Acid Research* online.

## Introduction

Molecular dynamics (MD) simulations are essential for understanding biomolecular systems at the atomic level. In the past, the high computational costs of these simulations severely limited their ability to achieve meaningful conformational exploration. However, this situation improved significantly with the introduction of Graphics Processing Units and algorithm advancements, which enabled researchers to study larger systems over longer timescales [1]. Despite the substantial progress in data production within the field [2, 3], it remains essential to develop novel and highly efficient post-processing methods to extract relevant insights from the simulated data.

Interaction fingerprint (IFP) analysis assists in elucidating the roles of key atoms or groups of atoms in molecular ensembles [4]. IFPs simplify the complexity of interatomic motions by translating them into the existence or absence of classical molecular interactions, including *π*-stacking, hydrogen bonds, and van der Waals contacts.

An interaction fingerprint is typically represented as a binary vector, where each entry signifies the presence or absence of a specific inter-atomic (or inter-residue) interaction. In this representation, a value of “1” indicates that a particular interaction is present, while a “0” means that the interaction does not occur. Generally, two interacting, non-overlapping selections of the molecular system are defined (e.g., ligand-protein, DNA-protein). However, intra-molecular selections (e.g., protein-protein, DNA-DNA) are also feasible at a stronger, usually prohibitive, performance penalty.

The applications of IFPs vary from tasks as simple as counting molecular interactions to more complex purposes, such as clustering conformational substates [5, 6]. IFPs can also be the starting point for intricate workflows to understand long-range non-covalent phenomena, such as allostery [7, 8, 9]. More Software Scope of application Molecular selections Reference recently, interaction fingerprints or closely related interaction-based descriptors have been adopted as input features for machine learning and deep learning models in structure-based drug discovery, including protein–ligand binding affinity prediction, pose ranking, and virtual screening [10, 11, 12, 13]. In this context, IFPs provide a compact and chemically interpretable representation of macromolecular interfaces that can be readily combined with neural network architectures, thereby enabling large-scale, dynamics-aware prediction tasks on simulation-derived conformational ensembles.

These opportunities are reinforced by the rapid development of FAIR repositories for MD trajectories. The Molecular Dynamics Data Bank (MDDB) aims to provide a unified European infrastructure for storing and reusing biosimulation data [14]. In parallel, MDRepo offers an open data warehouse for community-contributed protein MD simulations, explicitly designed to support large-scale downstream analyses and AI workflows [15]. More recently, DynaRepo has been introduced as a curated repository of macromolecular conformational dynamics, aggregating more than a millisecond of simulations for hundreds of complexes and single-chain systems, with precomputed analyses to facilitate data mining and dynamics-aware deep learning [16]. The emergence of such resources highlights the importance of fast and memory-efficient methods to compute interaction fingerprints at scale, so that rich interaction features can be extracted from ever-growing MD datasets.

Most software for detecting IFPs are designed to work with docking results (see Table 1). This task is generally less computationally demanding than analyzing MD simulations since it usually involves a much smaller number of poses and conformations. Despite this difference, the underlying algorithms remain similar. Many of them are equipped with visualization tools [17], although their reliance on web servers can complicate the analysis of large datasets. This dependency may also interfere with automated workflows, create challenges in online accessibility, or raise security concerns when managing sensitive data.

**Table 1.**
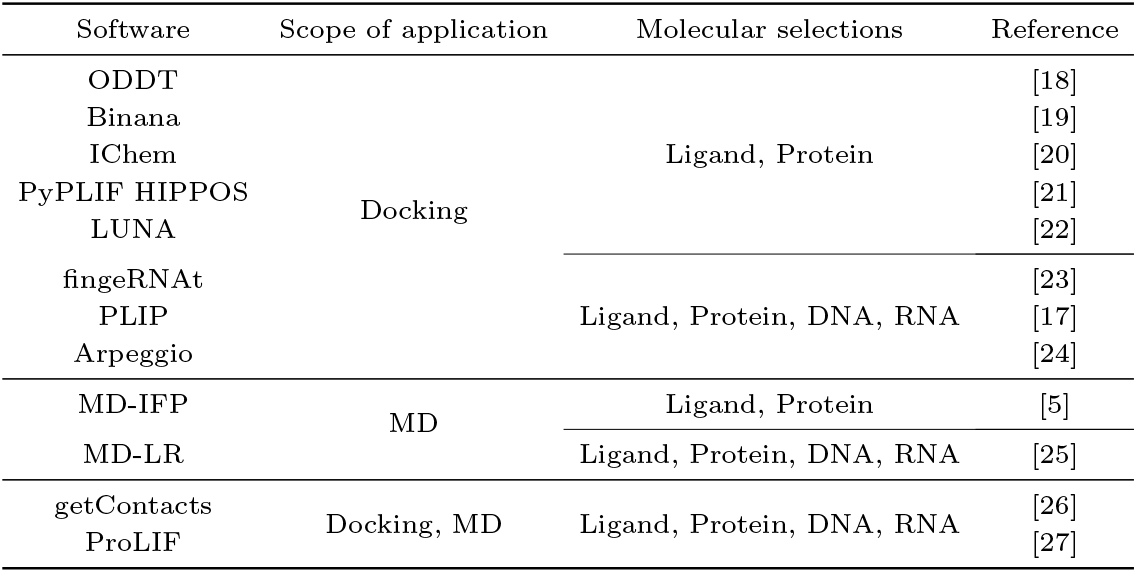
Scope of application of representative free software for computing IFPs.

In contrast to docking-focused tools, there are a few computational suites specifically designed for analyzing MD trajectories. Among the most frequently cited are getContacts [26], MD-IFP [5], MD-Ligand-Receptor [25], and notably ProLIF [27].

The getContacts scripts are based on VMD [28], and define atom selections with strict naming conventions that may not be compatible across various force fields. Additionally, several interactions are not defined for ligands, which reduces the versatility of this suite. MD-IFP can only treat ligand-protein or protein A-protein B systems, which prevents its applicability for detecting intra-molecular IFPs or simulations containing nucleic acids. On the other hand, MD-Ligand-Receptor, which executes PLIP (see Table 1) in parallel architectures, is restricted to working only with formats of the GROMACS [29] ecosystem. ProLIF addresses all the mentioned limitations, but much like the other methods, it is more effective when the molecular selections involve a limited number of atoms. Due to the inherent time and space complexities, all these tools face significant challenges when processing long trajectories of complex molecular systems or when tasked with computing intra-molecular IFPs, even for small systems.

From a practical perspective, the bottleneck of detecting IFPs consists of measuring interatomic distances (frequently) and angles (much less often), while adhering to specific geometric constraints. Introduced in 1975 by Bentley [30], k-dimensional trees (k-d trees) are highly effective for spatial searches in high-dimensional datasets. One of the primary operations performed with this data structure is neighbor searching, which can significantly reduce computational complexity from *O*(*N* ^2^) to *O*(*logN* ) in average cases or *O*(*N* ) in the worst case. This makes k-d trees a valuable and, to the best of our knowledge, under-exploited choice for IFP computing.

Here we present InterMap, a Python package designed to accelerate the calculation of IFPs in extensive MD trajectories of challenging systems. By utilizing k-d trees, InterMap significantly decreases the overall cost of distance calculations, which renders it particularly useful for detecting intra-molecular interactions. The package embraces Numba [31], an open-source just-in-time compiler that optimizes a subset of Python and NumPy [32] code into fast machine code. This choice improves computational speed and enables the execution of NumPy array expressions across multiple CPU cores without unnecessary memory overhead.

InterMap is designed to interface seamlessly with MDAnalysis, inheriting its ability to process various molecular formats and the rich selection syntax. MDAnalysis integration with RDKit [33] allows recovering molecular interaction patterns through SMiles Arbitrary Target Specification (SMARTS) [34], a regular expression language tailored for molecular systems. The well-defined nature of these patterns allows InterMap to operate independently of traditional residue or atomic naming conventions.

To facilitate data post-processing and minimize memory usage, InterMap efficiently compresses IFPs by encoding them as bit vectors using a bit array. This approach differs from the conventions of popular scientific Python libraries such as TensorFlow [35], NumPy [36], and SciPy [37], where the smallest storage alternative is an eightfold larger byte array. InterMap’s bit arrays are further compressed to prevent sparse fingerprints from consuming unnecessary resources. Several interactive visualizations are generated locally through a Shiny application equipped with convenient filters.

The package is freely available at https://github.com/Delta-Research-Team/intermap.git, while documentation and tutorials can be found at https://Delta-Research-Team.github.io/intermap/.

## Materials and Methods

### InterMap’s workflow

The general workflow followed by InterMap is illustrated in Figure 1. It can be launched either as a Python module or as a command-line interface (CLI). The CLI option provides a straightforward way for non-expert users to access the application, while the Python module is ideal for integrating InterMap into automated workflows without creating disruptions.

**Fig. 1.**
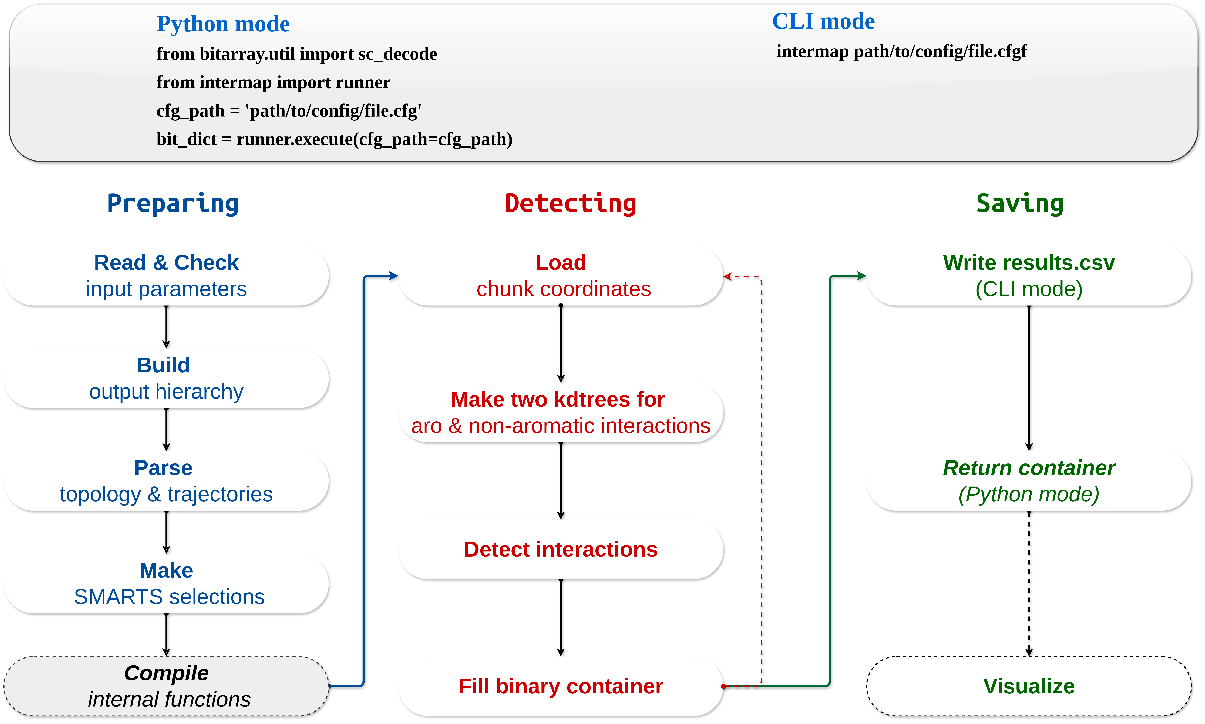
InterMap’s general workflow when launched either from a Python script (Python mode) or as a command line interface (CLI mode). The compilation step (in gray) only occurs the first time the software is run on a particular operating system

Once invoked, the first step in the application workflow is to parse the input and ensure that all parameters are correctly set. The software first establishes a directory hierarchy to organize the output files systematically. In addition to the IFPs binary results, the configuration file and a LOG file will be produced in the same output directory. It then loads the topology and trajectory data, which must have recognizable file extensions by MDAnalysis. Multiple replicas can be submitted in the same job.

The software analyzes two user-defined atom selections to compute the IFPs, utilizing SMARTS queries to identify the atomic indices that can engage in each available interaction (refer to Table 2). These selections must comply with the comprehensive selection syntax exposed by MDAnalysis. If it is the first time running InterMap on a particular operating system, all Numba-accelerated functions are compiled and cached to improve performance for subsequent runs.

**Table 2.**
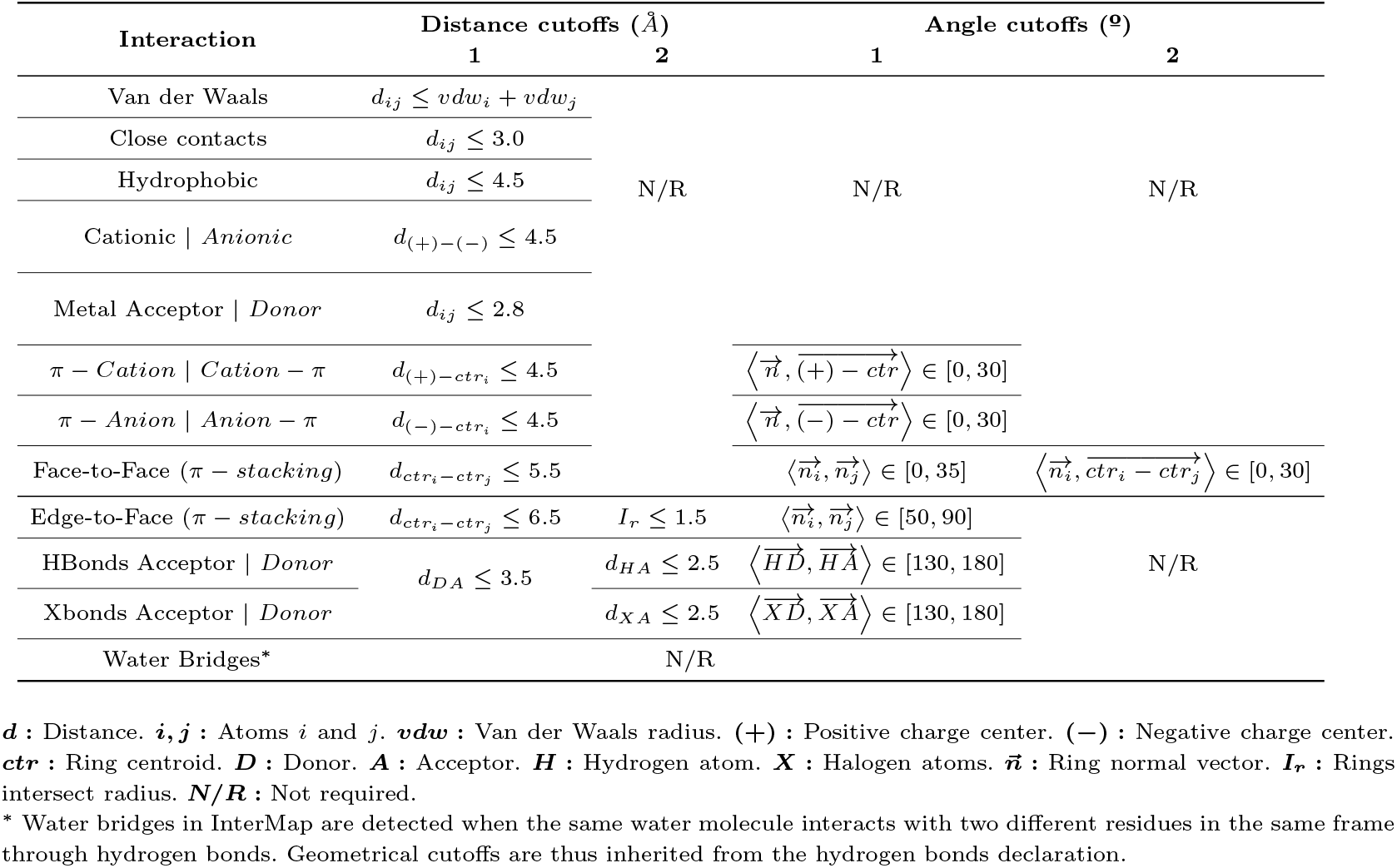
Type of interactions supported by InterMap and their default geometrical definitions.

The trajectory data is then processed in chunks of coordinates from which two k-d trees are constructed for each frame: one for aromatic and the other for non-aromatic interactions. This distinction is justified by the nature of the queries performed on the k-d trees. The query performance declines as the cutoff distance to identify neighbors increases or the number of neighbors involved rises. Non-aromatic interactions typically comprise more atoms but require a shorter cutoff distance. Conversely, the cutoff distance for aromatic interactions is usually higher, but fewer atoms are included in the calculations. As k-d trees are not valid data structures in the standard Python library or Numba ecosystems, they are constructed and queried using the numba-kdtree library [38].

Subsequently, the interaction detection phase takes place, during which a container of binary vectors is iteratively updated to capture information about which atoms are interacting, under what types of interactions, and in which frames these interactions occur. The frame assignment (either as “on” or “off”) is done inside a bit vector data structure provided by the bitarray project (https://github.com/ilanschnell/bitarray) that can assign a bit to each position. Finally, the software saves this container to the hard drive as a .*PICKLE* file (CLI mode) or returns a dictionary of compressed bit arrays (Python mode). Users can access interactive visualizations by loading the configuration file and the .*PICKLE* file into a local Shiny application, which enhances deriving insights from the data (see Section *InterVis: InterMap’s visualizations*).

Users can adjust the resolution or granularity of the computed IFPs by indicating whether they should be calculated at the inter-atomic (more detailed) or inter-residue (more condensed) level. It is also possible to specify a subset of interactions to compute. Furthermore, InterMap includes a feature to annotate parts of the molecular topology with personalized names, providing another way to label significant components that might not be adequately covered by conventional chain or segment identifiers. To this end, a file must be prepared in which each line corresponds to a distinct, non-overlapping set of atoms in the topology. When this option is used, the resulting outputs will include a column referencing these annotations for each detected interaction. This straightforward step can significantly enhance the ability to focus on key components of the interacting systems when processing or visualizing the IFPs data.

### Interaction definitions and SMARTS Patterns

InterMap can detect inter- and intra-molecular interactions for all biomolecules as long as they are recognizable by MDAnalysis. This includes small ligands, proteins, lipids, glycans, ions, solvents, DNA, and RNA. The available interactions and their default geometrical cutoffs are detailed in Table 2. Users can customize these internal defaults to to match their individual preferences.

Atoms or groups of atoms that may participate in these defined interactions are extracted from the input topology using queries based on the SMARTS patterns listed in Table 1 of the Supporting Information. Although the structure of these SMARTS patterns may initially appear cryptic, each term is clearly defined and can be decoded using the documentation from the Daylight project [34].

### Benchmark details

### MD Trajectories

We selected various MD trajectories as case studies to evaluate the computational performance of InterMap against the most relevant IFP software. The atomic composition of each system is detailed in Table 3, and their initial configurations are illustrated in Figure 2. These trajectories comprise systems of varying complexities, highlighting the diversity of biomolecular components commonly analyzed using IFPs tools. The original publications that describe the protocols for generating these trajectories are referenced in Table 3, along with the atomic selections used for computing IFPs.

**Table 3.**
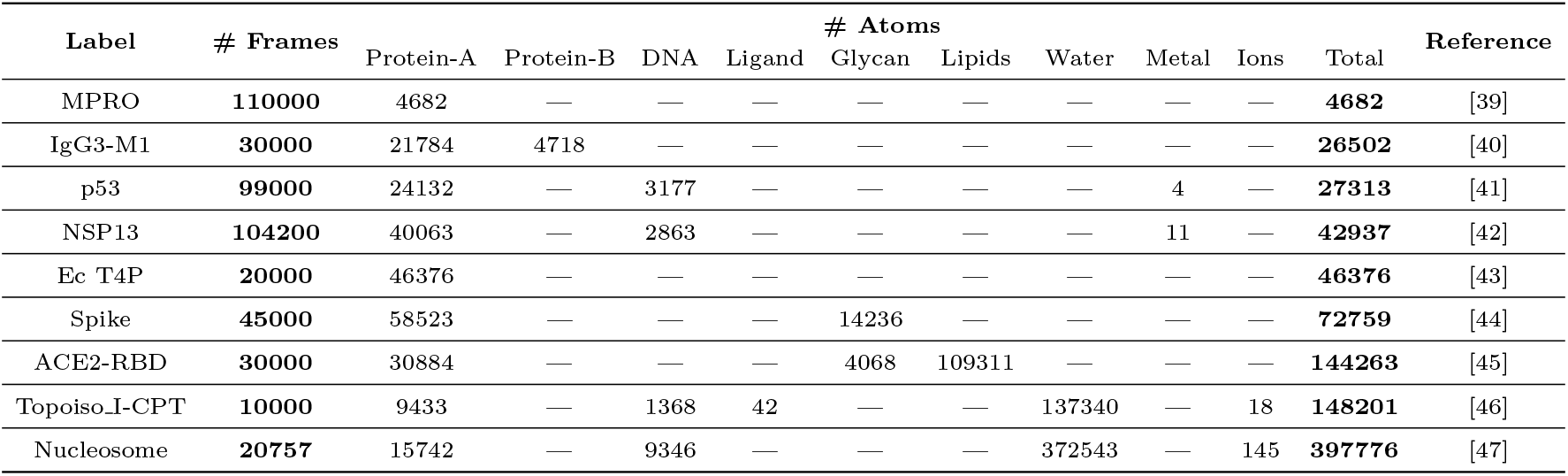
Molecular composition and number of frames of the analyzed MD trajectories.

| Label | # Frames | # Atoms |  |  |  |  |  |  |  |  |  | Reference |
| --- | --- | --- | --- | --- | --- | --- | --- | --- | --- | --- | --- | --- |
|  |  | Protein-A | Protein-B | DNA | Ligand | Glycan | Lipids | Water | Metal | Ions | Total |  |
| MPRO | **110000** | 4682 | — | — | — | — | — | — | — | — | **4682** | [39] |
| IgG3-M1 | **30000** | 21784 | 4718 | — | — | — | — | — | — | — | **26502** | [40] |
| p53 | **99000** | 24132 | — | 3177 | — | — | — | — | 4 | — | **27313** | [41] |
| NSP13 | **104200** | 40063 | — | 2863 | — | — | — | — | 11 | — | **42937** | [42] |
| Ec T4P | **20000** | 46376 | — | — | — | — | — | — | — | — | **46376** | [43] |
| Spike | **45000** | 58523 | — | — | — | 14236 | — | — | — | — | **72759** | [44] |
| ACE2-RBD | **30000** | 30884 | — | — | — | 4068 | 109311 | — | — | — | **144263** | [45] |
| Topoiso_I-CPT | **10000** | 9433 | — | 1368 | 42 | — | — | 137340 | — | 18 | **148201** | [46] |
| Nucleosome | **20757** | 15742 | — | 9346 | — | — | — | 372543 | — | 145 | **397776** | [47] |

**Fig. 2.**
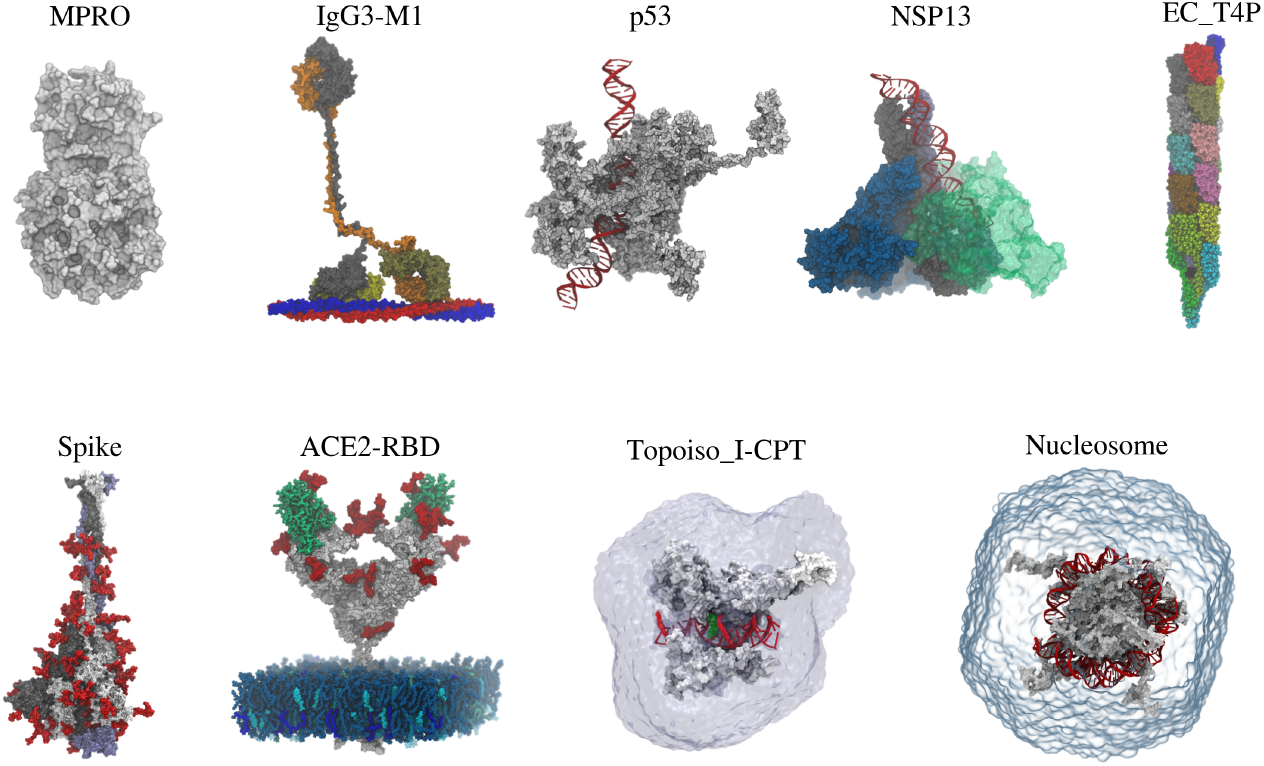
Initial conformations of the selected case studies. The Topoiso I–CPT and Nucleosome systems were processed with their full water boxes to evaluate InterMap’s performance on water–solute interactions.

**MPRO** refers to the main protease, a crucial enzyme in coronaviruses that mediates viral replication and transcription [39]. **IgG3-M1** is a complex formed between subclass 3 of Immunoglobulin (IgG3) and one of the most important virulence factors of the *Streptococcus pyogenes*, the M1 protein [40]. The tumor suppressor protein **p53** plays a vital role in regulating the cell cycle and preventing cancer [41]. **NSP13** is identified as SARS-CoV-2 non-structural protein 13, essential for viral replication [42]. The **Ec T4P** is the type IV Pilus of the *Escherichia coli*, that are large filaments at the surface of the bacteria with the ability to rapidly extend, retract and withstand strong forces [43]. The **Spike-open** refers to the SARS-CoV-2 spike glycoprotein trimer in the open state [44], while **ACE2-RBD** describes the complex of the spike receptor-binding domain (RBD) in its open state with angiotensin-converting enzyme 2 (ACE2) [45]. **Topoiso I-CPT** refers to the human DNA topoisomerase I in complex with camptothecin, a chemotherapeutic inhibitor that traps the enzyme in a cleavage complex with DNA [46]. Lastly, the **Nucleosome** refers to the first level of DNA compaction, featuring 147 base pairs wrapped around an octamer of histone proteins [47].

#### IFP software

InterMap was benchmarked against the following software: getContacts (https://getcontacts.github.io/, last commit on master branch: da14deb), MD-IFP v1.1, MD-LR v1.0, and ProLIF v2.0.0.

#### Environment

All software was executed on a hexa-core personal computer equipped with 64 GB of DDR4 RAM and an AMD Ryzen 5 processor operating at 3.2 GHz with hyperthreading support under Xubuntu 22.04 LTS. Wall-clock time and peak RAM usage were measured using the command “/usr/bin/time - v”. As the memory peaks reported by getContacts were not correctly captured by the previous command, we used an *in-house* script to gather all the child process initiated by this package (see *Section S2: getContacts RAM peak measure in the Supporting Information)*.

## Results and Discussion

### IFPs recovery: InterMap vs ProLIF

This section compares InterMap’s results against ProLIF, a widely used and actively maintained library for computing IFPs from MD trajectories. Both tools identify similar classes of non-covalent interactions under their default *residue* mode, while InterMap additionally supports a more detailed inter-atomic analysis via the *atom* mode.

Figure 3 provides a representative overview of the interactions observed between all residue pairs across 1000 equally spaced frames along the MPRO trajectory, where all intra-molecular interactions were considered. The data are presented as three boxplots per interaction type, each illustrating the distribution of the number of frames in which a given residue pair is found to interact. The green boxplots correspond to residue pairs detected by both InterMap and ProLIF, while the orange and red boxplots correspond to residue pairs detected exclusively by InterMap or ProLIF, respectively.

**Fig. 3.**
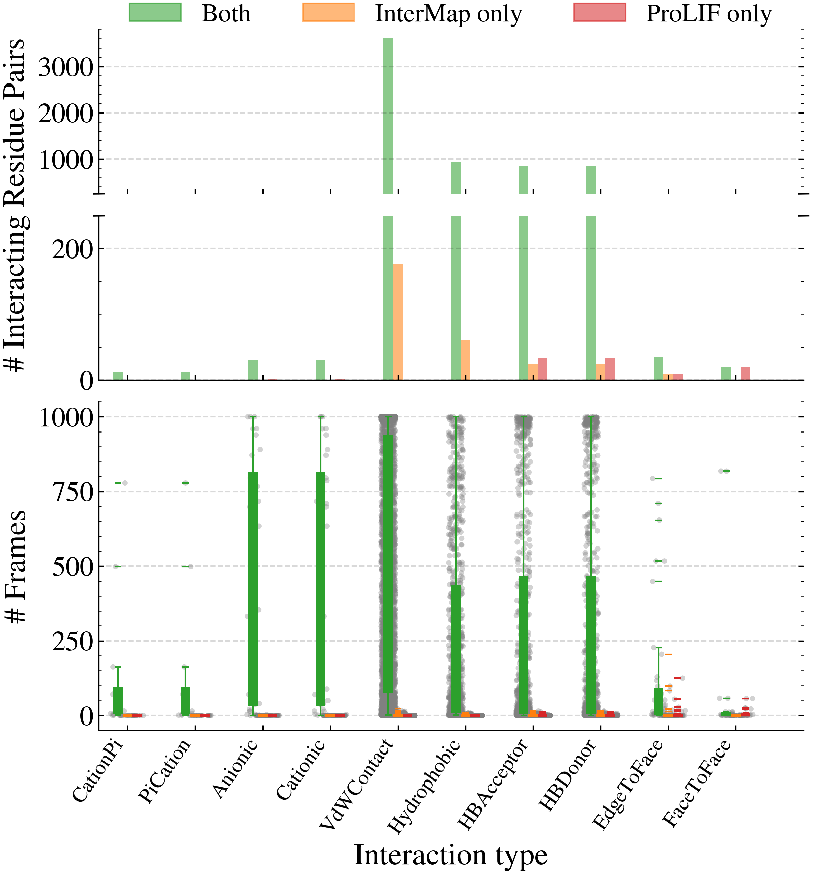
Number and frequency of inter-residue interactions detected by InterMap and ProLIF along 1000 equally-spaced frames of the MPRO trajectory. The lower panel shows, for each interaction type, the distribution of interaction lifetimes (number of frames in which a given inter-residue interaction is observed) as boxplots, with individual interaction frequencies overlaid as gray jittered points. For each interaction type, three distributions are reported: interactions detected by both methods (green), interactions detected only by InterMap (orange), and interactions detected only by ProLIF (red). The upper panels summarize, for the same interaction types and categories, the total number of distinct inter-residue pairs contributing to each distribution. A broken y-axis is used in the top block to simultaneously visualize low-count and high-count interaction types, with truncated bars in the lower top panel and corresponding full-height bars in the upper top panel.

In this case study, all residue pairs forming *π*–cation, cation–*π*, anionic or cationic interactions were detected by both tools in the same frames. Minor discrepancies were observed for the remaining interaction types, but a detailed inspection of the trajectories confirmed that their origin can be readily rationalized. For example, van der Waals contacts and hydrophobic interactions detected only by InterMap correspond to interatomic distances that lie within the chosen distance cutoff and are therefore chemically reasonable; these contacts are not reported by ProLIF (see Figure 1A,B in the Supporting Information for representative examples).

For hydrogen bonds, we identified a small subset of non-persistent interactions where ProLIF occasionally assigns peptide-bond nitrogen atoms as acceptors (Figure 1C in the Supporting Information). In such cases, the hydrogen attached to the genuine donor is oriented towards the amide nitrogen. This behaviour is inconsistent with the standard description of peptide hydrogen bonds, in which the amide nitrogen acts as a donor but not as an acceptor, as it lacks available lone pairs due to resonance stabilization of the peptide bond[48].

Both programs describe *π*–stacking interactions as a combination of edge-to-face and face-to-face orientations. Consequently, differences in how these orientations are defined can lead to variations in the classification of individual *π*–stacking contacts. These discrepancies likely arise from implementation details, such as the specific set of atoms used to construct aromatic ring normals. Nevertheless, visual inspection of these cases shows that both tools report chemically plausible *π*–stacking patterns, and that the majority of persistent interactions are identified in the same frames.

In summary, the observed discrepancies between InterMap and ProLIF are minor and do not affect the scientific conclusions derived from IFP analyses performed with either tool.

### Performance comparison of IFP software

We benchmarked InterMap against ProLIF, getContacts, and MD-LR using a diverse set of nine MD trajectories. These trajectories included various systems: protein–protein, protein–DNA, protein–ligand, and large solvated complexes. For each software, we analyzed wall-clock time, peak memory usage, and the total number of detected interactions, both at the residue and atom levels when applicable. The results are summarized in Table 4.

**Table 4.**
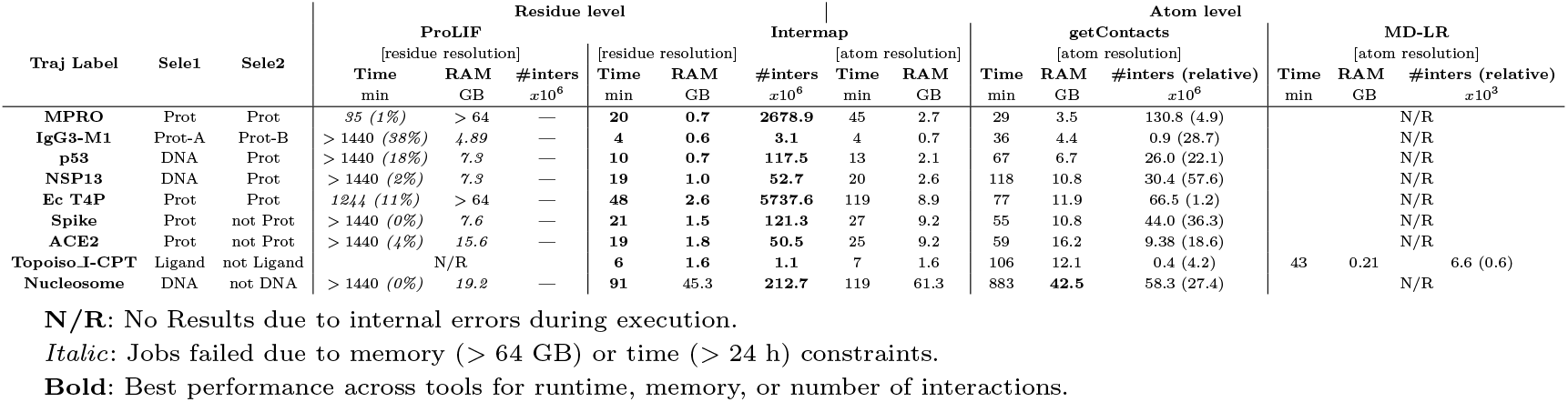
Performance comparison of IFP software for processing MD trajectories. The number of detected interactions is reported in its absolute value (#inters) and the relative percentage with respect to interMap (relative).

In “residue” mode, InterMap represents interactions at the residue level, resulting in faster execution times and lower memory consumption. Conversely, “atom” mode provides a higher resolution analysis by examining interactions at the atomic level, which requires more computational resources but also offers more detailed insights. Its dual-mode capability enables users to balance analytical depth with computational efficiency based on their specific research needs.

In “residue” resolution, InterMap consistently achieved the fastest execution times across all analyzed trajectories, ranging from 3 minutes for protein-protein interactions (IgG3-M1) to 91 minutes for the Nucleosome complex while maintaining remarkably stable memory consumption between 0.6 and 2.6 GB for most systems. Notably, InterMap successfully completed analysis of all trajectories, detecting substantially more interactions than competing methods while maintaining predictable scaling with system complexity.

According to its original design, which is focused on studying ligand dissociation rather than IFP analysis as the primary objective, MD-IFP (not shown in Table 4) has significant limitations when computing IFPs. Its operation is constrained to trajectories that include a small ligand and a protein or two distinct proteins, which limits its effectiveness for general IFP analyses and prevents it from processing any of our benchmarked trajectories.

MD-LR presents several key limitations in biomolecular interaction analysis. Due to its architectural design, the software cannot process nucleic acids as independent ligands, significantly limiting its application in DNA/RNA-protein interaction studies. Additionally, its direct derivation from the PLIP framework means MD-LR inherits the inability to compute intra-molecular IFPs, preventing analysis of internal conformational dynamics of macromolecules. A further constraint requires all ligand atoms to be assigned to a single molecular group, which must be explicitly specified during input configuration. This structural limitation effectively narrows MD-LR’s practical utility to small molecule ligands, making it unsuitable for larger or more complex molecular systems. MD-LR is also restricted to the input formats of the GROMACS ecosystem. These limitations considerable shrinks its application scope, forcing us to include the trajectory provided by the software’s authors in order to compare against InterMap (TopoisoI-CPT in Table 4). This use case took 43 minutes and 0.2 GB with MD-LR, against 7 minutes and 1.6 GB with InterMap. It is worth noting that MD-LR detected only 0.6% of the interactions recovered by InterMap.

getContacts displayed highly variable temporal performance, with execution times ranging from 29 minutes in MPRO to 882 minutes for the Nucleosome. While this tool completed all analyses, its performance deteriorated with system size or when computing intra-molecular IFPs, requiring 4-10 fold longer execution times than InterMap for comparable systems. Most critically, getContacts consistently detected orders of magnitude fewer interactions than InterMap. For instance, it identified only 130.8 million interactions in MPRO, accounting for just 4.9% of what InterMap detects, and 66.49 million in Ec T4P, which represents a mere 1.2% of InterMap’s calculations. These figures highlight a fundamental limitation in the definition of interactions in getContacts.

The most striking differences are observed when analyzing the performance of ProLIF, which faces severe computational bottlenecks. In the case of the MPRO trajectory, for example, ProLIF took 35 minutes to complete just 1% of the analysis, reaching a memory consumption over 64 GB. Based on a linear extrapolation, the complete analysis would require approximately 3500 minutes. Similar results were observed in other trajectories: IgG3-M1 (3789 minutes), p53 (8000 minutes), and NSP13 (72000 minutes). In contrast, InterMap completed these same analyses in only 3, 10, and 15 minutes, respectively, representing performance improvements of two and three orders of magnitude. Worse, in the Spike and Nucleosome case studies, ProLIF could not even start the simulation after one day of loading.

InterMap’s memory efficiency stands as a distinctive advantage, maintaining consistently low RAM consumption across all systems while simultaneously detecting the highest number of interactions.

getContacts, except for the Nucleosome case, consistently required more RAM than InterMap for equivalent analyses, ranging from 3.5 to 42.5 GB. This memory overhead becomes particularly troubling when considering its reduced interaction detection capability. For instance, getContacts utilized 11.9 GB to identify 66.49 million interactions in Ec T4P, while InterMap calculated 86 times more interactions using just 8.9 GB. Additionally, in the Nucleosome case, although getContacts consumed slightly less RAM than InterMap, it only detected about 27% of the interactions reported by InterMap.

ProLIF exhibited the most severe memory limitations, repeatedly exceeding the 64 GB threshold and crashing due to memory exhaustion. The software’s memory consumption makes it unsuitable for medium to large molecular systems. Even for the smallest trajectory (MPRO), ProLIF consumed the maximum available 64 GB, representing a 91-fold higher memory requirement than InterMap’s 0.7 GB.

Overall, the benchmark shows that InterMap combines generality (“residue” and “atom” resolutions, compatibility with heterogeneous systems), two to three orders of magnitude faster wall-clock times, and a low, predictable memory footprint while recovering substantially more interactions than existing tools.

#### InterMap’s scalability trade-offs

While InterMap is significantly faster and more memory-efficient than other IFP software designed for MD, it does exhibit specific bottlenecks and limitations that influence its scalability for handling complicated tasks. This section addresses some identified issues, translating them into general user recommendations.

We assessed time and RAM peak usage across the two available resolution modes, *atom* and *residue* (see Section *InterMap’s workflow*), over 50000 frames of the MPRO trajectory (see Figure 4). The MPRO case study represents one of the most demanding scenarios in computing IFPs: recovering intra-molecular interactions. Compared to interfacial analyses, this requires examining far more possible distance and angle relationships within a single macromolecule. To speed up our analysis, we limited it to a specific portion of the trajectory; however, the overall trends would remain consistent. Specifically, we examined the performance impact of variations in the number of processors engaged to run InterMap and the specified chunk size (the number of frames loaded simultaneously into RAM).

**Fig. 4.**
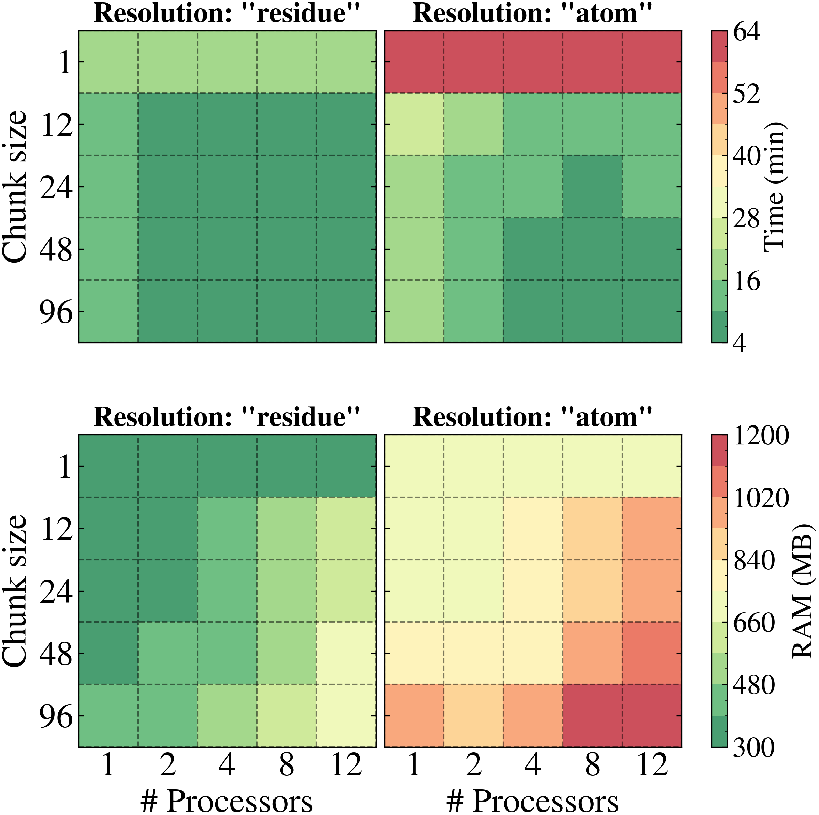
Impact of chunk size and processor count on InterMap’s performance while processing the MPRO trajectory. Both resolution modes (*atom* and *residue*) are illustrated.

InterMap execution comprises three primary steps that are repeated throughout a run, as illustrated in Figure 1-center: (i) loading N frames from disk, (ii) detecting IFPs for each frame in parallel, and (iii) storing the detected IFPs as bitarrays in an appropriate data structure. Among these, only the second step, which involves detecting IFPs, is parallelized. As a result, the first and third steps become bottlenecks in the overall execution.

Using a chunk size of a single frame consistently leads to poor runtime performance, regardless of the number of processors engaged (see Figure 4, top panel). This outcome is expected since the detection functions are parallelized. Also, it is clear that opting for *residue* resolution significantly outperforms *atom* resolution regarding speed. This improvement occurs because, in *residue* resolution, detected interactions at the atomic level are aggregated per residue, resulting in fewer entries to store. When the chunk size is enlarged, a substantial speedup is observed, which is linked to fewer disk read operations required. However, further increases beyond a reduced number of frames yield only mild or negligible improvements in runtime. This trend can again be attributed to the fact that storing, rather than detecting IFPs or loading frames, remains the slowest step in the workflow.

Increasing the number of processors can also reduce runtime, but this improvement is not linear and exhibits a conservative trend. InterMap does not benefit from an excessively high processor count, even when parallelization is applied because the subsequent storage step remains computationally intensive. To optimize InterMap runtime, users should ideally set the chunk size to match the number of processors. However, as discussed below, using too many processors may increase memory consumption without providing proportional performance gains. The increase of chunk size has a pronounced effect on RAM consumption, which depends on the number of coordinates per frame (Figure 4, bottom panel). For systems larger than MPRO, increasing the chunk size may significantly increase memory usage. In the MPRO trajectory, time performance reaches a plateau beyond a particular chunk size, while RAM consumption continues growing, highlighting the need for a balanced configuration. Additional processor count also increases RAM demands due to the concurrent existence of multiple copies of different data structures. When memory availability is constrained, users are advised to reduce the number of processors and the chunk size to complete a run that otherwise may have crashed due to a lack of memory resources.

### InterVis: InterMap’s visualizations

InterMap offers robust functionality through its locally hosted web application, InterVis (see Figure 5), which provides a comprehensive framework for getting general insights from the IFPs detected during an MD simulation.

**Fig. 5.**
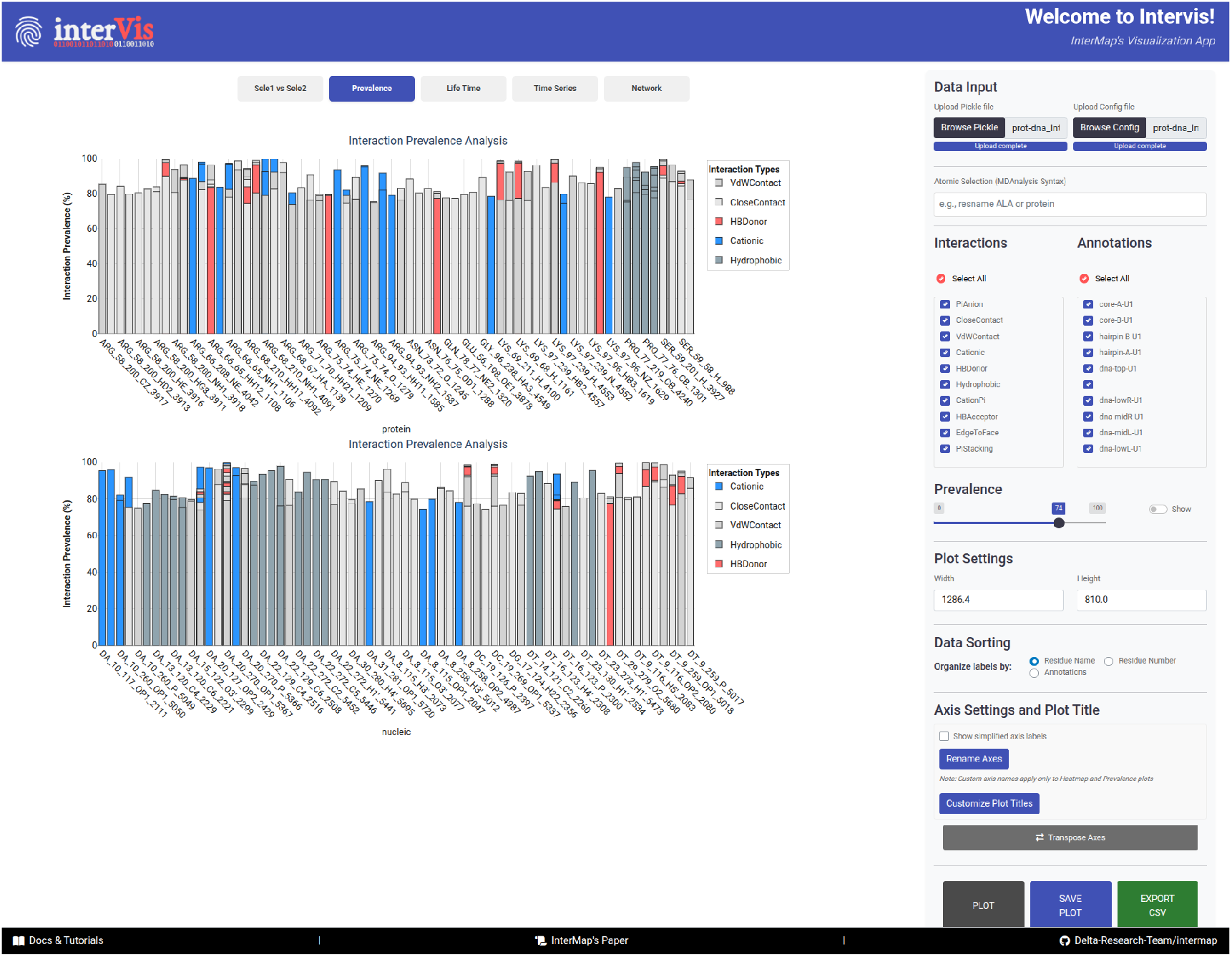
An example of an Intervis session is illustrated. The right gray panel enables users to make precise selections, which generate interactive plots displayed in the left panel. The screenshot displays the plots created for the “Prevalence” analysis, which indicates the percentage of the trajectory in which a particular interaction is active. The top and bottom panels highlight each defined interacting partner.

The only inputs required for InterVis are the configuration file used during the InterMap run and the corresponding *ICKLE* output file. With this information, users can efficiently select and filter raw data, generating interactive plots that provide in-depth insights into the system’s interaction patterns. Users can refine the plotted information using several criteria: (i) the type of interactions, (ii) atomic annotations (if included in the configuration file; see Section: *InterMap’s workflow*), (iii) a prevalence cutoff, or (iv) a proper MDAnalysis atomic selection (enabled by the internal mapping of the input topology to the .*PICKLE* atomic labels).

After applying these filters, InterVis generates five types of interactive plots, each displayed in a separate tab. The first tab presents a 2D heatmap illustrating the interactions established between both selections used during the IFP computation. The second tab features two bar plots that show the prevalence of interactions for Selection 1 and Selection 2. The third tab includes a box plot depicting the distribution of interaction lifetimes, where each value corresponds to the number of consecutive frames in which a particular interaction was detected. The fourth tab offers a time series analysis of interaction presence across frames. Finally, an interactive network visualization represents residues as nodes and displays edges between interacting residues, color-coded by interaction type, with line width proportional to the prevalence of interactions.

These plots can be exported as images in .png format. Additionally, filtered data can be exported in .csv format, facilitating further analysis outside InterVis or the creation of handcrafted external plots. Representative images and detailed explanations for each plot type are available in *Section S4: Visualizations* of the Supporting Information, further clarifying the functionalities and analytical capabilities offered by InterVis. The powerful combination of versatile filtering options and interactive plots makes InterVis an effective tool for quickly gaining exploratory insights into the expression and evolution of interactions within simulations.

## Conclusion

In this study, we present InterMap, a Python package designed for the rapid and memory-efficient detection of IFPs in large-scale MD trajectories that significantly outperforms existing software. InterMap uses several optimization strategies, including k-d trees for spatial neighbor searches to reduce the computational complexity of distance queries significantly, Numba just-in-time compilation to accelerate critical loops and facilitate multicore execution without substantial memory overhead, and bit array compression to represent IFPs as compact binary vectors. Additionally, it integrates with MDAnalysis and utilizes SMARTS patterns, offering users flexible selection options across various biomolecular formats. An interactive Shiny-based local web viewer, InterVis, further enhances exploratory analysis capabilities.

Our benchmarks reveal that InterMap can achieve up to 99% reductions in both runtime and peak memory usage compared to established alternatives such as ProLIF, getContacts, MD-IFP, and MD-LR, across diverse case studies involving small proteins, large glycoprotein complexes, and nucleosome systems. With both Command Line Interface and Python API, InterMap supports both manual analyses and automated pipelines, allowing researchers to customize parameters such as interaction definitions, atom versus residue resolution, chunk size, and parallelism.

We anticipate that InterMap will become a standard tool in MD post-processing toolkits, facilitating more profound insights into critical biomolecular mechanisms, including allostery, ligand binding, and conformational dynamics of complex systems.

## Author contributions statement

R.G-A. and Y.K conceived the study and developed the main ideas. E.F. and R.G.A. wrote the initial draft of the manuscript and programmed the software. Y.K. and E.B. contributed to the overall conceptual framework and performed the data collection. F.D. analyzed the results, contributed to the discussion, and managed funding and resources for the research. All authors contributed to the revisions and approved the final version of the manuscript.

## Acknowledgments

The data used in this work was generated with computing HPC and storage resources by GENCI at IDRIS on the supercomputer Jean Zay (grants 2022-A0130713814 and 2023-A0150714577), and by the EXPLOR center hosted by the University de Lorraine (grant 2019CPMXX0983). R.G.-A.’s contribution to this work was supported by ANR grant ANR-22-CE11-0029-01.

---

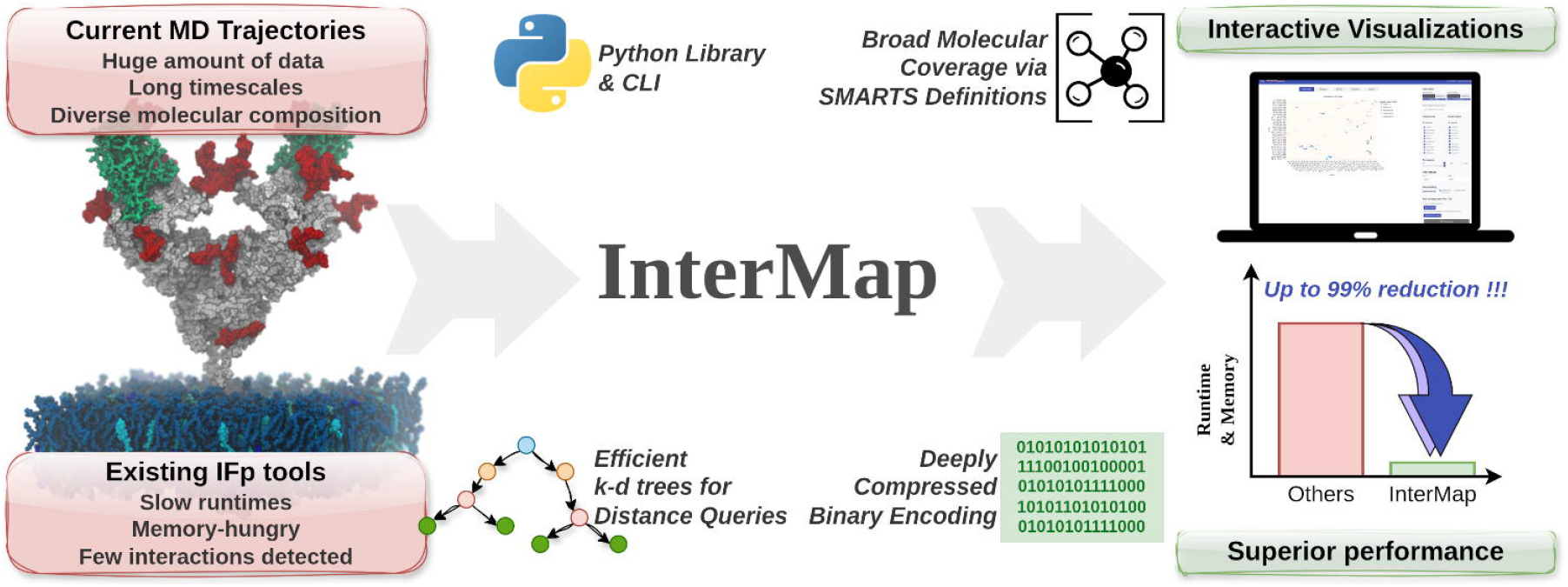

## References

1. Scott A. Hollingsworth and Ron O. Dror. Molecular dynamics simulation for all. 99(6):1129–1143. Publisher: Elsevier.

2. Francesco Madeddu, Jessica Di Martino, Michele Pieroni, Davide Del Buono, Paolo Bottoni, Lorenzo Botta, Tiziana Castrignanò, and Raffaele Saladino. Molecular docking and dynamics simulation revealed the potential inhibitory activity of new drugs against human topoisomerase i receptor. 23(23):14652.

3. Davide Pirolli, Benedetta Righino, Chiara Camponeschi, Francesco Ria, Gabriele Di Sante, and Maria Cristina De Rosa. Virtual screening and molecular dynamics simulations provide insight into repurposing drugs against SARS-CoV-2 variants spike protein/ACE2 interface. 13(1):1494.

4. Zhan Deng, Claudio Chuaqui, and Juswinder Singh*. Structural interaction fingerprint (SIFt): A novel method for analyzing three-dimensional proteinligand binding interactions. Archive Location: world Publisher: American Chemical Society.

5. Daria B. Kokh, Bernd Doser, Stefan Richter, Fabian Ormersbach, Xingyi Cheng, and Rebecca C. Wade. A workflow for exploring ligand dissociation from a macromolecule: Efficient random acceleration molecular dynamics simulation and interaction fingerprint analysis of ligand trajectories. 153(12):125102.

6. Parker Ladd Bremer, Danna De Boer, Walter Alvarado, Xavier Martinez, and Eric J. Sorin. Overcoming the heuristic nature of k-means clustering: Identification and characterization of binding modes from simulations of molecular recognition complexes. 60(6):3081–3092. Publisher: American Chemical Society.

7. Mitul Srivastava, Lovika Mittal, Anita Kumari, and Shailendra Asthana. Molecular dynamics simulations reveal the interaction fingerprint of remdesivir triphosphate pivotal in allosteric regulation of SARS-CoV-2 RdRp. 8. Publisher: Frontiers.

8. Divine Mensah Sedzro, Mukhtar Oluwaseun Idris, Olanrewaju Ayodeji Durojaye, Abeeb Abiodun Yekeen, Adeola Abraham Fadahunsi, and Suleiman Oluwaseun Alakanse. Identifying potential p53-MDM2 interaction antagonists: An integrated approach of pharmacophore-based virtual screening, interaction fingerprinting, MD simulation and DFT studies. 7(39):e202202380. eprint: https://onlinelibrary.wiley.com/doi/pdf/10.1002/slct.202202380.

9. Sneha Bheemireddy, Roy González Alemán, Emmanuelle Bignon, and Yasaman Karami. Communication pathway analysis within protein-nucleic acid complexes. bioRxiv, pages 2025–02, 2025.

10. Zoe Li, Ruili Huang, Menghang Xia, Tucker A. Patterson, and Huixiao Hong. Fingerprinting Interactions between Proteins and Ligands for Facilitating Machine Learning in Drug Discovery. Biomolecules, 14(1):72, January 2024.

11. Christian Fellinger, Thomas Seidel, Benjamin Merget, Klaus-Juergen Schleifer, and Thierry Langer. GRADE and X-GRADE: Unveiling Novel Protein–Ligand Interaction Fingerprints Based on GRAIL Scores. Journal of Chemical Information and Modeling, 65(5):2456–2475, March 2025. Publisher: American Chemical Society.

12. Manuel S. Sellner, Markus A. Lill, and Martin Smieško. Quality Matters: Deep Learning-Based Analysis of Protein-Ligand Interactions with Focus on Avoiding Bias, November 2023. Pages: 2023.11.13.566916 Section: New Results.

13. Jeremy Wohlwend, Gabriele Corso, Saro Passaro, Mateo Reveiz, Ken Leidal, Wojtek Swiderski, Tally Portnoi, Itamar Chinn, Jacob Silterra, Tommi Jaakkola, and Regina Barzilay. Boltz-1 democratizing biomolecular interaction modeling. Pages: 2024.11.19.624167 Section: New Results.

14. Molecular Dynamics Data Bank. The European Repository for Biosimulation Data | MDDB | Project | Fact Sheet | HORIZON.

15. Amitava Roy, Ethan Ward, Illyoung Choi, Michele Cosi, Tony Edgin, Travis S Hughes, Md Shafayet Islam, Asif M Khan, Aakash Kolekar, Mariah Rayl, Isaac Robinson, Paul Sarando, Edwin Skidmore, Tyson L Swetnam, Mariah Wall, Zhuoyun Xu, Michelle L Yung, Nirav Merchant, and Travis J Wheeler. MDRepo—an open data warehouse for community-contributed molecular dynamics simulations of proteins. Nucleic Acids Research, 53(D1):D477–D486, January 2025.

16. Omid Mokhtari, Emmanuelle Bignon, Hamed Khakzad, and Yasaman Karami. DynaRepo: the repository of macromolecular conformational dynamics. Nucleic Acids Research, page gkaf1130, November 2025.

17. Melissa F. Adasme, Katja L. Linnemann, Sarah Naomi Bolz, Florian Kaiser, Sebastian Salentin, V. Joachim Haupt, and Michael Schroeder. PLIP 2021: expanding the scope of the protein–ligand interaction profiler to DNA and RNA. 49:W530–W534. Publisher: Oxford Academic.

18. Maciej Wójcikowski, Piotr Zielenkiewicz, and Pawel Siedlecki. Open drug discovery toolkit (ODDT): a new open-source player in the drug discovery field. 7(1):26.

19. Jade Young, Neerja Garikipati, and Jacob D. Durrant. BINANA 2: Characterizing Receptor/Ligand Interactions in Python and JavaScript. Journal of Chemical Information and Modeling, 62(4):753–760, February 2022. Publisher: American Chemical Society.

20. Franck Da Silva, Jeremy Desaphy, and Didier Rognan. IChem: A versatile toolkit for detecting, comparing, and predicting protein-ligand interactions. 13(6):507–510.

21. Enade P. Istyastono, Muhammad Radifar, Nunung Yuniarti, Vivitri D. Prasasty, and Sudi Mungkasi. PyPLIF HIPPOS: A molecular interaction fingerprinting tool for docking results of AutoDock vina and PLANTS. 60(8):3697–3702. Publisher: American Chemical Society.

22. Alexandre V. Fassio, Laura Shub, Luca Ponzoni, Jessica McKinley, Matthew J. O’Meara, Rafaela S. Ferreira, Michael J. Keiser, and Raquel C. de Melo Minardi. Prioritizing virtual screening with interpretable interaction fingerprints. 62(18):4300–4318.

23. Natalia A. Szulc, Zuzanna Mackiewicz, Janusz M. Bujnicki, and Filip Stefaniak. fingeRNAt—a novel tool for high-throughput analysis of nucleic acid-ligand interactions. 18(6):e1009783. Publisher: Public Library of Science.

24. Harry C Jubb, Alicia P Higueruelo, Bernardo Ochoa-Montaño, Will R Pitt, David B Ascher, and Tom L Blundell. Arpeggio: A web server for calculating and visualising interatomic interactions in protein structures. 429(3):365–371.

25. Michele Pieroni, Francesco Madeddu, Jessica Di Martino, Manuel Arcieri, Valerio Parisi, Paolo Bottoni, and Tiziana Castrignanò. MD–ligand–receptor: A high-performance computing tool for characterizing ligand–receptor binding interactions in molecular dynamics trajectories. 24(14):11671. Number: 14 Publisher: Multidisciplinary Digital Publishing Institute.

26. A. J. Venkatakrishnan, Rasmus Fonseca, Anthony K. Ma, Scott A. Hollingsworth, Augustine Chemparathy, Daniel Hilger, Albert J. Kooistra, Ramin Ahmari, M. Madan Babu, Brian K. Kobilka, and Ron O. Dror. Uncovering patterns of atomic interactions in static and dynamic structures of proteins. Pages: 840694 Section: New Results.

27. Cédric Bouysset and Sébastien Fiorucci. ProLIF: a library to encode molecular interactions as fingerprints. 13(1):72.

28. William Humphrey, Andrew Dalke, and Klaus Schulten. VMD – Visual Molecular Dynamics. Journal of Molecular Graphics, 14:33–38, 1996.

29. Mark James Abraham, Teemu Murtola, Roland Schulz, Szilárd Páll, Jeremy C. Smith, Berk Hess, and Erik Lindahl. GROMACS: High performance molecular simulations through multi-level parallelism from laptops to supercomputers. SoftwareX, 1-2:19–25, September 2015.

30. Jon Louis Bentley. Multidimensional binary search trees used for associative searching. Communications of the ACM, 18(9):509–517, 1975.

31. Siu Kwan Lam, stuartarchibald, Antoine Pitrou, Mark Florisson, Graham Markall, Stan Seibert, Emergency Self-Construct, Todd A. Anderson, Guilherme Leobas, rjenc29, Michael Collison, Kaustubh, luk-f a, Jay Bourque, Aaron Meurer, Travis E. Oliphant, Nick Riasanovsky, Michael Wang, densmirn, KrisMinchev, Andre Masella, njwhite, Ethan Pronovost, Eric Wieser, Ehsan Totoni, Stefan Seefeld, Hernan Grecco, Pearu Peterson, Isaac Virshup, and Matty G. numba/numba: Numba 0.61.0, January 2025.

32. Charles R. Harris, K. Jarrod Millman, Stéfan J. van der Walt, Ralf Gommers, Pauli Virtanen, David Cournapeau, Eric Wieser, Julian Taylor, Sebastian Berg, Nathaniel J. Smith, Robert Kern, Matti Picus, Stephan Hoyer, Marten H. van Kerkwijk, Matthew Brett, Allan Haldane, Jaime Fernández del Río, Mark Wiebe, Pearu Peterson, Pierre Gérard-Marchant, Kevin Sheppard, Tyler Reddy, Warren Weckesser, Hameer Abbasi, Christoph Gohlke, and Travis E. Oliphant. Array programming with NumPy. 585(7825):357–362. Publisher: Nature Publishing Group.

33. RDKit.

34. Daylight theory: SMARTS - a language for describing molecular patterns.

35. Martín Abadi, Ashish Agarwal, Paul Barham, Eugene Brevdo, Zhifeng Chen, Craig Citro, Greg S. Corrado, Andy Davis, Jeffrey Dean, Matthieu Devin, Sanjay Ghemawat, Ian Goodfellow, Andrew Harp, Geoffrey Irving, Michael Isard, Yangqing Jia, Rafal Jozefowicz, Lukasz Kaiser, Manjunath Kudlur, Josh Levenberg, Dandelion Mané, Rajat Monga, Sherry Moore, Derek Murray, Chris Olah, Mike Schuster, Jonathon Shlens, Benoit Steiner, Ilya Sutskever, Kunal Talwar, Paul Tucker, Vincent Vanhoucke, Vijay Vasudevan, Fernanda Viégas, Oriol Vinyals, Pete Warden, Martin Wattenberg, Martin Wicke, Yuan Yu, and Xiaoqiang Zheng. TensorFlow: Large-scale machine learning on heterogeneous systems, 2015. Software available from tensorflow.org.

36. Charles R. Harris, K. Jarrod Millman, Stéfan J. van der Walt, Ralf Gommers, Pauli Virtanen, David Cournapeau, Eric Wieser, Julian Taylor, Sebastian Berg, Nathaniel J. Smith, Robert Kern, Matti Picus, Stephan Hoyer, Marten H. van Kerkwijk, Matthew Brett, Allan Haldane, Jaime Fernández del Río, Mark Wiebe, Pearu Peterson, Pierre Gérard-Marchant, Kevin Sheppard, Tyler Reddy, Warren Weckesser, Hameer Abbasi, Christoph Gohlke, and Travis E. Oliphant. Array programming with NumPy. Nature, 585(7825):357–362, September 2020.

37. Pauli Virtanen, Ralf Gommers, Travis E. Oliphant, Matt Haberland, Tyler Reddy, David Cournapeau, Evgeni Burovski, Pearu Peterson, Warren Weckesser, Jonathan Bright, Stéfan J. van der Walt, Matthew Brett, Joshua Wilson, K. Jarrod Millman, Nikolay Mayorov, Andrew R. J. Nelson, Eric Jones, Robert Kern, Eric Larson, C J Carey, İlhan Polat, Yu Feng, Eric W. Moore, Jake VanderPlas, Denis Laxalde, Josef Perktold, Robert Cimrman, Ian Henriksen, E. A. Quintero, Charles R. Harris, Anne M. Archibald, Antônio H. Ribeiro, Fabian Pedregosa, Paul van Mulbregt, and SciPy 1.0 Contributors. SciPy 1.0: Fundamental Algorithms for Scientific Computing in Python. Nature Methods, 17:261–272, 2020.

38. Felix Igelbrink. mortacious/numba-kdtree. original-date: 2021-10-01T11:26:08Z.

39. Seyed Arad Moghadasi, Emmanuel Heilmann, Ahmed Magdy Khalil, Christina Nnabuife, Fiona L. Kearns, Chengjin Ye, Sofia N. Moraes, Francesco Costacurta, Morgan A. Esler, Hideki Aihara, Dorothee von Laer, Luis Martinez-Sobrido, Timothy Palzkill, Rommie E. Amaro, and Reuben S. Harris. Transmissible SARS-CoV-2 variants with resistance to clinical protease inhibitors. Science Advances, 9(13):eade8778, March 2023. Publisher: American Association for the Advancement of Science.

40. Arman Izadi, Yasaman Karami, Eleni Bratanis, Sebastian Wrighton, Hamed Khakzad, Maria Nyblom, Berit Olofsson, Lotta Happonen, D. Tang, Martin Sundwall, Magdalena Godzwon, Yashuan Chao, Alejandro Gomez Toledo, Tobias Schmidt, Mats Ohlin, Michael Nilges, Johan Malmström, Wael Bahnan, Oonagh Shannon, Lars Malmström, and Pontus Nordenfelt. The hinge-engineered IgG1-IgG3 hybrid subclass IgGh47 potently enhances Fc-mediated function of anti-streptococcal and SARS-CoV-2 antibodies. Nature Communications, 15(1):3600, April 2024. Publisher: Nature Publishing Group.

41. Ö Demir, P. U. Ieong, and R. E. Amaro. Full-length p53 tetramer bound to DNA and its quaternary dynamics. Oncogene, 36(10):1451–1460, March 2017.

42. D. E. Shaw Research. Molecular dynamics simulations related to sars-cov-2. D. E. Shaw Research Technical Data, 2020.

43. Yasaman Karami, Aracelys López-Castilla, Andrea Ori, Jenny-Lee Thomassin, Benjamin Bardiaux, Therese Malliavin, Nadia Izadi-Pruneyre, Olivera Francetic, and Michael Nilges. Computational and biochemical analysis of type iv pilus dynamics and stability. Structure, 29(12):1397–1409, 2021.

44. Lorenzo Casalino, Zied Gaieb, Jory A. Goldsmith, Christy K. Hjorth, Abigail C. Dommer, Aoife M. Harbison, Carl A. Fogarty, Emilia P. Barros, Bryn C. Taylor, Jason S. McLellan, Elisa Fadda, and Rommie E. Amaro. Beyond Shielding: The Roles of Glycans in the SARS-CoV-2 Spike Protein. ACS Central Science, September 2020. Publisher: American Chemical Society.

45. Emilia P. Barros, Lorenzo Casalino, Zied Gaieb, Abigail C. Dommer, Yuzhang Wang, Lucy Fallon, Lauren Raguette, Kellon Belfon, Carlos Simmerling, and Rommie E. Amaro. The flexibility of ACE2 in the context of SARS-CoV-2 infection. Biophysical Journal, 120(6):1072–1084, March 2021. Publisher: Elsevier.

46. Michele Pieroni. MD-ligand-receptor input trajectory test case. June 2023.

47. Yasaman Karami and Emmanuelle Bignon. Cysteine hyperoxidation rewires communication pathways in the nucleosome and destabilizes the dyad. Computational and Structural Biotechnology Journal, 23:1387–1396, 2024.

48. E.N. Baker and R.E. Hubbard. Hydrogen bonding in globular proteins. Progress in Biophysics and Molecular Biology, 44(2):97–179, 1984.

