## Supplementary material for "InterMap: Accelerated Detection of Interaction Fingerprints on Large-Scale Molecular Ensembles": Suplementary information

*Nucleic Acid Research,* 2025, pp. 1–4

**doi: DOI HERE**
Advance Access Publication Date: Day Month Year
Supporting Information for:

SUPPORTING INFORMATION FOR:

### InterMap: Accelerated Detection of Interaction Fingerprints on Large-Scale Molecular Ensembles

Ernesto Fajardo-Díaz,[1] Emmanuelle Bignon,[2] François Dehez,[2] Yasaman Karami[3,*] and Roy González-Alemán[2,3*]

[1]Université Paris Cité, 75013, Paris, France, [2]Université de Lorraine, CNRS, UMR 7019, LPCT, F-54000 Nancy, France and [3]Université de Lorraine, CNRS, Inria, LORIA, F-54000 Nancy, France
FOR PUBLISHER ONLY Received on Date Month Year; revised on Date Month Year; accepted on Date Month Year

#### S1: SMART patterns defined in InterMap

Table 1 contains the SMART patterns defined in InterMap and a brief description of their meaning. These patterns are taken from ProLIF, and were inspired by Pharmit (https://sourceforge.net/p/pharmit/code/ci/master/tree/src/pharmarec.cpp) or RDKit (https://github.com/rdkit/rdkit/blob/master/Data/BaseFeatures.fdef).

#### S2: getContacts RAM peak measure

The RAM peak of all software benchmarked against InterMap was recorded using the Linux command "**/usr/bin/time -v**". However, the getContacts script starts several child processes, and the aforementioned command does not fully capture their resource consumption. To ensure a fair comparison, we used the script shown in **Algorithm 1** to compute the RAM peak as the sum of all those child processes.

**Algorithm 1** Monitor Process Memory Usage

```
get_all_pids() {
    local _pid=$1
    local _pids="$_pid"
    for child in $(pgrep -P $_pid); do
        _pids+",$(get_all_pids $child)"
    done
    echo $_pids
}

while kill -0 $_pid 2>/dev/null; do
    _pids=$(get_all_pids $_pid | tr "," " ")
    current_mem=$(ps -o rss= -p $_pids | awk '{sum+=$1} END {print sum}')
    [$current_mem -gt $max_mem] && max_mem=$current_mem
    sleep 0.5
done
```

© The Author 2025. Published by Oxford University Press. All rights reserved. For permissions, please

**Table 1.** Summary of SMARTS patterns used in InterMap and their plain English interpretation.

| Interaction Type | SMARTS Pattern | Plain English Description |
|---|---|---|
| Anions | [-{1-},$(O=[C,S,P]-[O-])] | Atoms with a formal negative charge, including deprotonated carboxylate, sulfonate, and phosphate oxygens (O=[C,S,P]–O$^-$). |
| Cations | [+{1-},$([NX3&!$([NX3]-O)]-[C]=[NX3+])] | Atoms with a formal +1 charge, including iminium-like systems: trigonal N–C=N$^+$, excluding N directly bound to O. |
| Hydrogen Bond Acceptors (hb_acc) | [$([N&!$([NX3]-*=[O,N,P,S])&!$([NX3]-[a])&!$([Nv4+1])&!$(N=C(-[C,N])-N)]),$([n+0&!X3&!$([n&r5]:[n+&r5])]),$([O&!$([OX2](C)C=O)&!$(O(~a)~a)&!$(O=N-*)&!$([O-]-N=O)]),$([o+0]),$([F&$(F-[#6])&!$(F-[#6][F,Cl,Br,I])])] | Neutral N (non-amide, non-nitro, non-quaternary), neutral aromatic N, non-ester/non-nitro O, neutral aromatic O, and C-bound F (not polyhalogenated) that can accept H-bonds. |
| Hydrogen Bond Donors (hb_don) | [$([O,S,#7;+0]),$([Nv4+1]),$([n+]c[nH])]-[H] | Neutral O, S, N, protonated tetravalent N, and specific aromatic cationic N with an explicit H that can donate H-bonds. |
| Hydrophobic (hydroph) | [c,s,Br,I,S&H0&v2,$([C&R0;$([CH0](=*)=*),$([CH1](=*)-[!#1]),$([CH2](-[!#1])-[!#1])]),$([C;$([CH0](=*)(-[!#1])-[!#1]),$([CH1](-![#1])(-[!#1])-[!#1])]),$([C&D4!R](-[CH3])(-[CH3])-[CH3]);!$([#6]~[#7,#8,#9]);+0] | Aromatic C, aromatic S, Br, I, S (v2, no H), and selected aliphatic C (incl. tert-butyl–like) not adjacent to N/O/F and neutral, representing hydrophobic moieties. |
| Metal Acceptors (metal_acc) | [O,#7&!$([nX3])&!$([NX3]-*=[!#6])&!$([NX3]-[a])&!$([NX4]),-{1-};!+{1-}] | O and non-aromatic, non-amide, non-quaternary N, plus negatively charged atoms (not +1), that can coordinate metals. |
| Metal Donors (metal_don) | [Ca,Cd,Co,Cu,Fe,Mg,Mn,Ni,Zn] | Metal ions: Ca, Cd, Co, Cu, Fe, Mg, Mn, Ni, and Zn as coordination centers. |
| 5-Membered Rings (rings5) | [a&r5]1:[a&r5]:[a&r5]:[a&r5]:[a&r5]:1 | Aromatic 5-membered rings with all five atoms aromatic and in the same ring. |
| 6-Membered Rings (rings6) | [a&r6]1:[a&r6]:[a&r6]:[a&r6]:[a&r6]:[a&r6]:1 | Aromatic 6-membered rings with six aromatic atoms in a single ring. |
| Water | [O&H2] | Explicit water molecules. |
| Halogen Bond Acceptors (xb_acc) | [#7,#8,P,S,Se,Te,a;!+{1-}]! #[*] | Electron-rich N, O, P, S, Se, Te, or aromatic atoms, not positively charged, that can accept halogen bonds (excluding triple-bonded atoms). |
| Halogen Bond Donors (xb_don) | [#6,#7,Si,F,Cl,Br,I]-[Cl,Br,I,At] | Halogens (Cl, Br, I, At) bound to C, N, Si, or F, acting as halogen-bond donors via a $\sigma$-hole. |

#### S3: InterMap–ProLIF interaction mismatches

The Figure 1 shows some representative examples of interactions detected either by InterMap or ProLIF only.

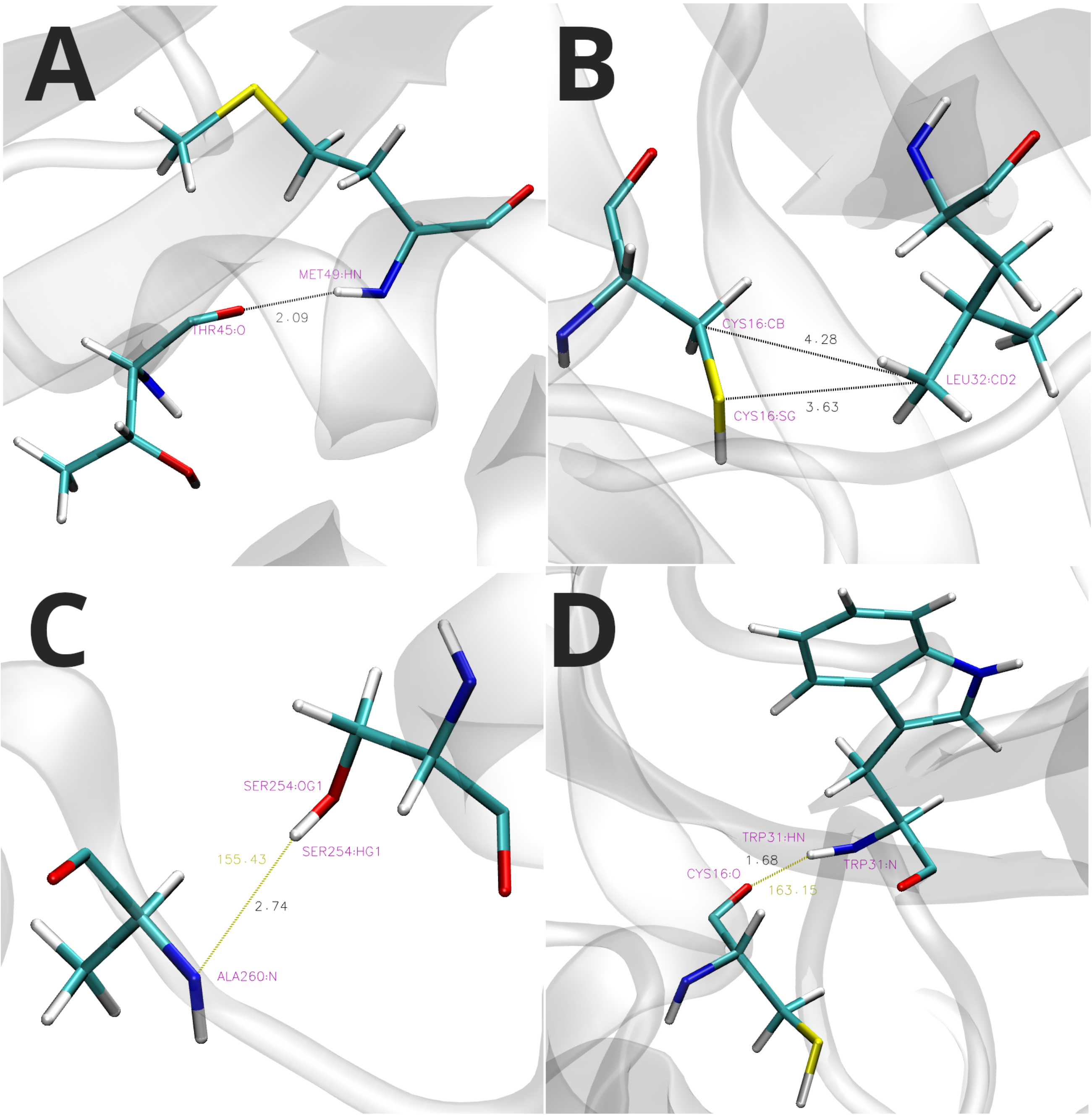


**Fig. 1.** Representative examples of mismatches between InterMap and ProLIF when recovering interactions. Black numbers refer to distances while olive ones refers to donor-hydrogen-acceptor angles. A: van der Waals contact detected by InterMap and not reported by ProLIF, B: Hydrophobic contacts detected by InterMap and not reported by ProLIF, C: Amide nitrogen reported as hydrogen bond acceptor by ProLIF and not detected by InterMap, and D: Hydrogen bond acceptor in cysteine detected by InterMap and not reported by ProLIF.

#### S4: Visualizations

The InterVis interface consists of an interactive, multi-tab analysis panel designed to explore and understand molecular interaction patterns from Molecular Dynamics simulations. Each visualization is organized into separate tabs for greater clarity and specific analytical purposes.

The "Selection 1 vs Selection 2" tab (Figure 2) displays a comprehensive interaction matrix that overviews all interactions between the selected molecular entities. Each cell in the matrix represents a specific type of interaction between two entities, which may correspond to individual atoms or entire residues, as defined by the user in the InterMap configuration file. The cell color indicates the interaction class, and a numerical value may optionally show its prevalence. When hovering over a cell, detailed information is displayed, including entity identifiers, annotations, interaction type, and prevalence percentage. This visualization is an excellent starting point for identifying general patterns and critical interaction points.

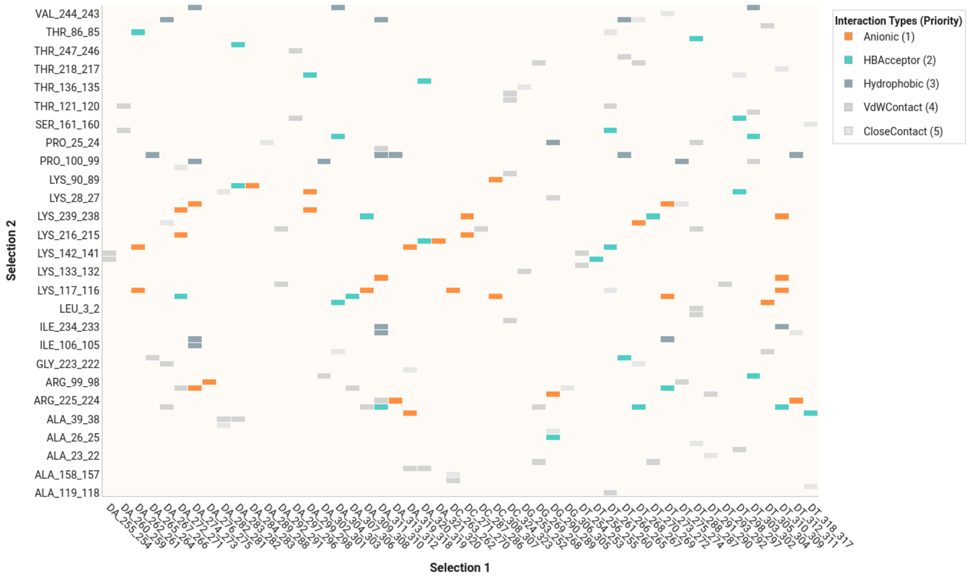


**Fig. 2.** Interaction matrix between entities from selections Selection 1 and Selection 2. Color indicates the interaction type; optional numbers show prevalence.

The "Prevalence" tab (Figure 3) contains two complementary bar charts: one focused on Selection 1 and another on Selection 2. Each chart shows the prevalence of different interaction types between an element from one selection and all its partners in the other. Interactions are grouped by the elements of Selection 1 (top plot) or Selection 2 (bottom plot), allowing exploration of individual contributions to the interaction network. Each bar is color-coded by interaction type, and hovering reveals further details such as interacting elements, prevalence percentage, and functional annotations.

The "Lifetime" tab (Figure 4) presents boxplots representing the duration of interactions between selected pairs throughout the simulation. Each violin shows the distribution of lifetimes (in frames) for a specific interaction type between a pair, with the width indicating the density of duration values. Box elements inside each violin reflect the median and interquartile range, while outliers are shown as individual points. A consistent color scheme identifies each interaction type. Hovering over violins or outliers reveals the interaction pair, type, prevalence, and specific frame ranges, facilitating the comparison between stable and transient interactions.

The "Time Series" tab (Figure 5) presents a dynamic view of how interactions evolve during the simulation. In the main panel, each marker represents an interaction event between a pair of elements at a specific frame, color-coded by interaction type. The horizontal histogram above shows the number of interactions per frame, while the vertical bar chart on the right summarizes the overall prevalence of each interacting pair. Hovering over any marker provides detailed metadata including frame number, interacting elements, interaction type, and prevalence. This visualization helps identify temporal patterns and potential coordinated events among molecular interactions.

The "Network" tab (Figure 6) provides an interactive graph representation of molecular interactions. In this visualization, nodes represent elements from Selection 1 and Selection 2, color-coded for distinction (e.g., blue for Selection 1 and red for Selection 2), while edges indicate interactions between nodes. The edges are styled such that their color represents the interaction type and their width reflects prevalence. Users can rearrange nodes by dragging, zoom in and out, and pan across the graph to explore spatial relationships. Hovering over edges or nodes reveals detailed metadata, including interacting elements, interaction type and prevalence. This visualization is particularly useful for identifying highly connected components, dominant interaction types, and structural organization within the molecular network.

Each tab has been optimized for interactive exploration, with hover-enabled details, customizable views, and consistent color coding across all graphs. This unified design facilitates comparisons and cross-referencing between different aspects of the interaction analysis.

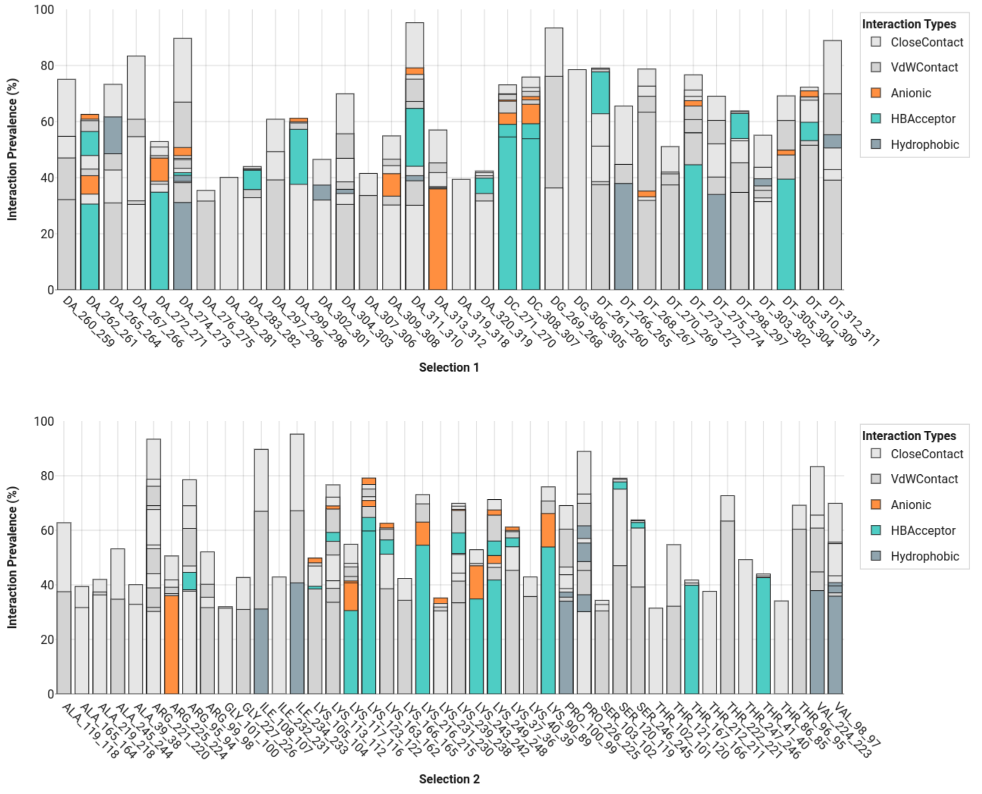


**Fig. 3.** Prevalence of interaction types grouped by elements of Selection 1 (top) and Selection 2 (bottom). Color denotes interaction type.

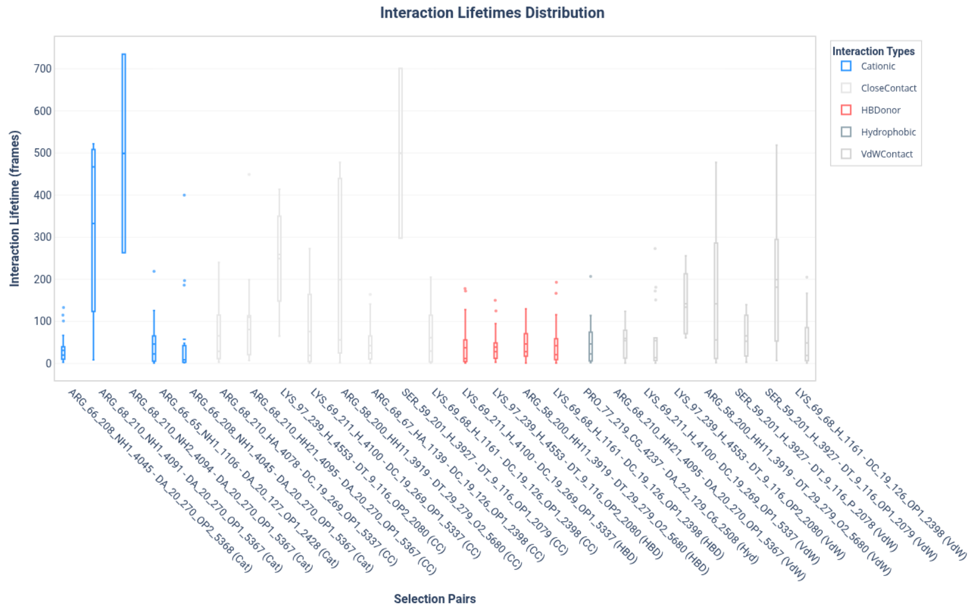


**Fig. 4.** Boxplots showing the lifetime distribution of interactions between entity pairs. Width indicates density; boxes show medians and ranges. Colors denote interaction type.

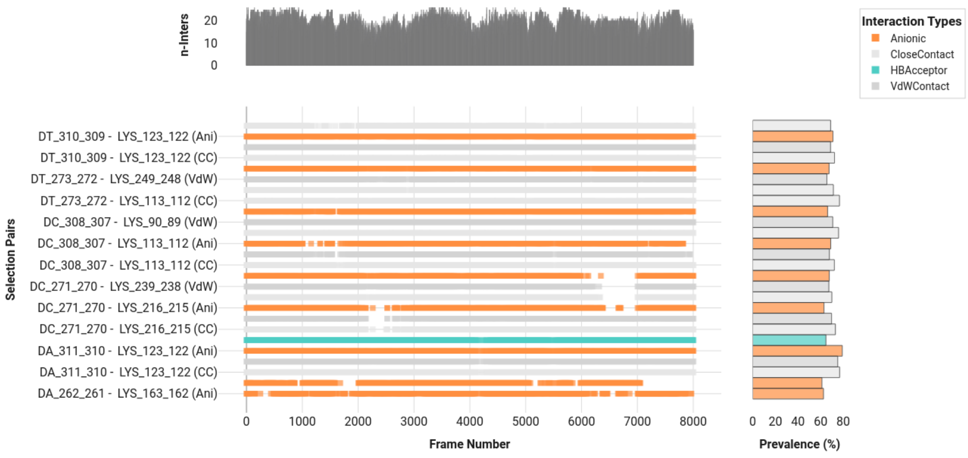


**Fig. 5.** Time-resolved visualization of interactions with temporal distribution (top), interaction events (main), and prevalence summary (right). Color encodes interaction type.

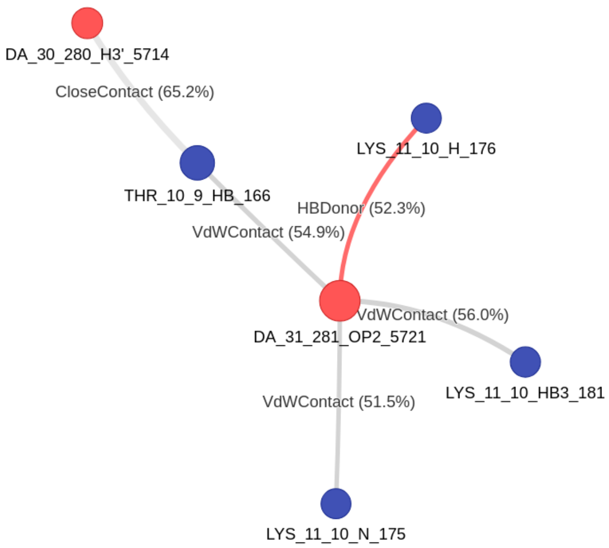


**Fig. 6.** Network-based visualization of molecular interactions, where nodes represent elements and edges encode interaction type (color) and prevalence (width).
